# EPOP and MTF2 Activate PRC2 Activity through DNA-sequence specificity

**DOI:** 10.1101/2025.09.25.678634

**Authors:** Jeffrey Granat, Sanxiong Liu, Luis Popoca, Ozgur Oksuz, Danny Reinberg

## Abstract

Polycomb Repressive Complex 2 (PRC2) facilitates the formation of facultative heterochromatin, instrumental to tissue specific gene expression. PRC2 catalyzes tri-methylation of lysine 27 of histone H3 (H3K27me3), which is targeted for chromatin compaction by PRC1. Importantly, PRC2-associated cofactors regulate its distinct activities, as in the case of MTF2 and JARID2 that direct PRC2 to specific chromatin nucleation sites based on preferred DNA-binding motifs. Here, we investigated EPOP whose role in regulating PRC2 was not well-defined. We find that both EPOP and MTF2 stimulate PRC2 histone methyltransferase (HMT) activity *in vitro*. Unlike MTF2, EPOP is ineffectual in PRC2 chromatin recruitment as evidenced by an EED-rescue system *in vivo*, but promotes H3K27me3 deposition *de novo* in cooperation with MTF2 and JARID2. Binding assays using reconstituted dinucleosome substrates revealed that similar to MTF2, EPOP promotes PRC2 chromatin-binding activity in a distinct DNA-sequence dependent manner (GCN-rich and GA-rich, respectively). Thus, EPOP and MTF2 in conjunction with JARID2 foster PRC2-mediated HMT activity at chromatin sites comprising cofactor-preferred DNA-binding sequences during the formation of H3K27me3-chromatin domains.

## Introduction

The accurate establishment and maintenance of developmentally-regulated patterns of transcriptional silencing by the Polycomb Repressive Complex 2 (PRC2) are closely tied to the regulation of its catalytic activity and recruitment to chromatin^1^. In *Drosophila*, repressor proteins target and recruit PRC2 to precise Polycomb Response Elements (PREs) to initiate the establishment of chromatin domains enriched in the product of PRC2 catalysis: tri-methylated lysine 27 of histone H3 (H3K27me3). These cis-acting regulatory DNA elements are necessary to sustain the transmission of H3K27me3-decorated chromatin throughout development, as their absence after the initial deposition of H3K27me3 results in a dilution of H3K27me3 and ultimately, transcriptional derepression^2–4^. In mammalian cells, however, there are only two documented cases of genetic elements that function as PREs^5,6^, and until recently, a general paradigm was lacking to explain genome-wide binding patterns of PRC2.

Recent work has provided insight into this quandary by defining the initial sites within chromatin to which PRC2 is recruited in mouse embryonic stem cells (mESCs)^7^. These “nucleation sites” are discrete foci that serve as hubs for the *de novo* recruitment of PRC2. Subsequently, H3K27me3 is spread distally both in *cis* and in *trans* through three-dimensional spatial interactions to establish broad domains of H3K27me3-repressive chromatin. Importantly, nucleation sites are found within CpG islands (CGIs) that are highly enriched in GCN-(N being any nucleotide) and GA-tandem repeats, relative to all other CGIs in the genome. Significantly, these tandem repeats bear resemblance to previously defined DNA binding motifs of PRC2 accessory subunits, such as MTF2 and JARID2^8,9^, which presumably target PRC2 to chromatin. Indeed, it was shown that nucleation sites are highly enriched in these cofactors, and cells lacking both MTF2 and JARID2 fail to accurately coordinate *de novo* recruitment of PRC2 to chromatin^7^. This scenario points to a mechanistic link between PRC2 cofactors and GCN- and GA-tandem repeats in coordinating PRC2 recruitment and deposition of H3K27me3. Multiple studies in the past several years have demonstrated that cofactors play overlapping but also non-redundant roles in establishing H3K27me3-domains^10–12^, suggesting that these accessory subunits exhibit site-specific functions. Yet, our understanding of the unique contributions of each cofactor to the establishment and maintenance of H3K27me3-domains remains vague.

Although PRC2 cofactors like MTF2 and JARID2 clearly aid PRC2 in establishing H3K27me3-repressive domains^7^, the functional role of another such accessory factor, EPOP, is not clear. Despite its enrichment at PRC2 nucleation sites^7^, previous findings attribute EPOP with a negative regulatory function through its hindrance of both PRC2 recruitment and PRC2 histone methyltransferase (HMT) activity^13,14^. Therefore, a central inquiry addressed in this study concerns the role of EPOP in the initial recruitment of PRC2 to chromatin and in the formation of *de novo* H3K27me3-domains. We also provide insight into the mechanistic basis by which EPOP and MTF2 might stimulate PRC2 HMT activity at chromatin site-specific regions.

## Results

### EPOP Stimulates PRC2 Histone Methyltransferase Activity in vitro

PRC2 accessory factors that are highly enriched at nucleation sites, such as MTF2 and JARID2, are not only crucial for the initial recruitment of PRC2 to chromatin^7^ but also have potent stimulatory effects on PRC2 HMT activity^9,15–19^. Despite its elevated enrichment at PRC2 nucleation sites^7^, EPOP is believed to inhibit both PRC2 recruitment to chromatin and subsequent deposition of H3K27me3. However, the studies supporting this conclusion involved knockdown experiments in cells, but not direct biochemical evidence^13,14^. Instead, we sought to determine the effect of EPOP on PRC2 HMT activity by performing *in vitro* HMT assays on oligonucleosome arrays using PRC2 complexes generated recombinantly. Both MTF2 and JARID2 robustly stimulated PRC2 HMT activity relative to the PRC2 core complex (PRC2-core; **Figure 1**). Although this effect had been previously demonstrated for JARID2^15,17^, this is the first demonstration that MTF2 directly stimulates PRC2 HMT activity *in vitro*, to our knowledge. Strikingly, the effect of EPOP on PRC2 HMT activity is more robust (**Figure 1**). Thus, contrary to previous speculation^13,14^ and similar to JARID2, MTF2 and EPOP directly stimulated PRC2 HMT activity *in vitro*.

**Figure 1.**
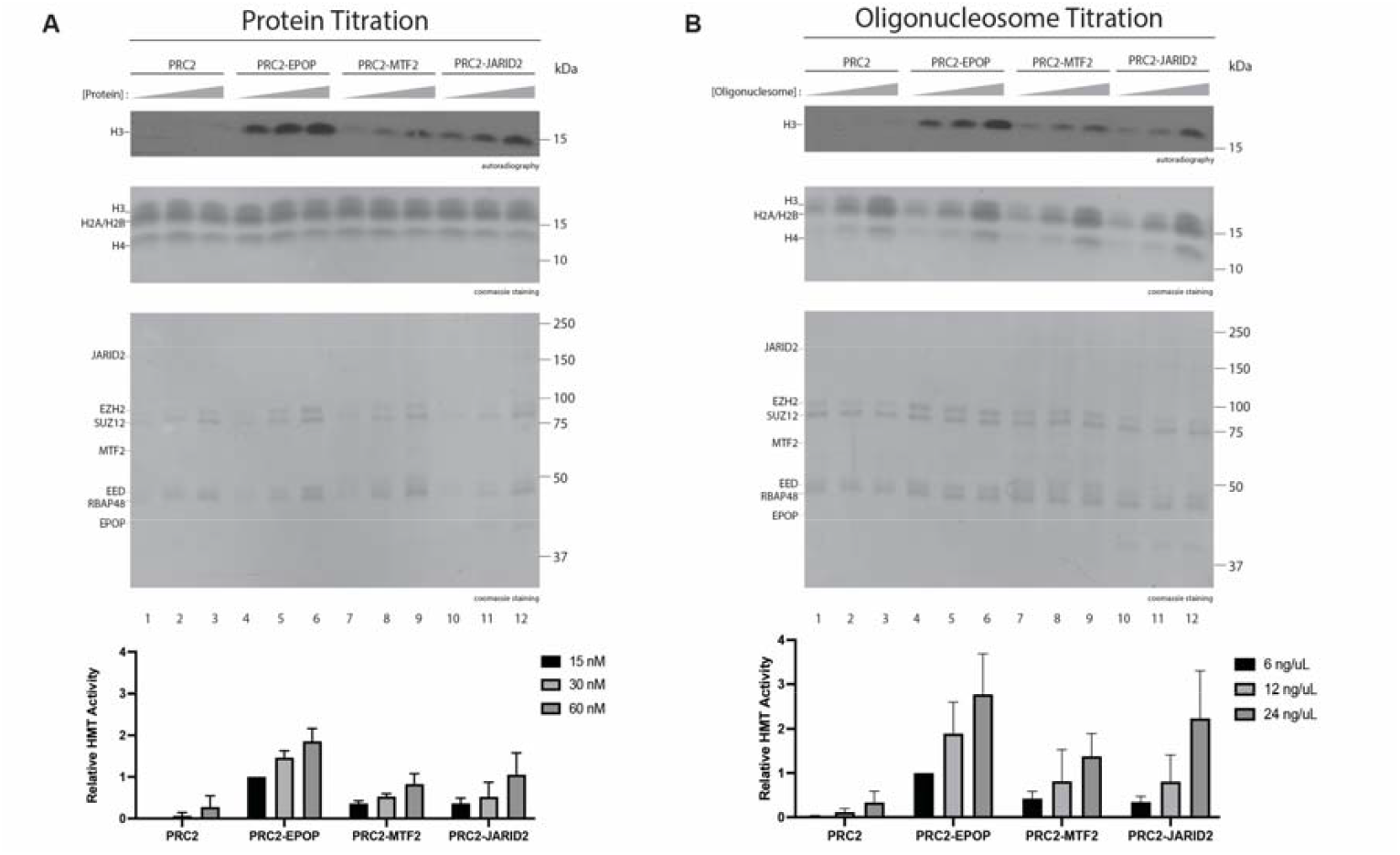
The HMT activity of PRC2 is promoted by EPOP, MTF2, and JARID2. (A, B) HMT assays (30 min incubation) performed with increasing amounts (15, 30, or 60 nM) of PRC2, PRC2-EPOP, PRC2-MTF2, and PRC2-JARID2 in the presence of 12x-oligonucleosome arrays (25 ng/uL (A) or with 60 nm of each PRC2 complex with increasing amounts of oligonucleosome arrays (6, 12, or 24 ng/uL) (B). H3 methylation levels were gauged by autoradiography (top images). Histones present in oligonucleosomes (middle images) and subunits of PRC2 complexes (bottom images) are shown by Coomassie blue staining of SDS-PAGE gels. Bottom panels: Quantification of relative amounts of ^3^H-SAM incorporated into H3, normalized to PRC2-EPOP (15 nM) (A) or PRC2-EPOP with 6 ng/uL oligonucleosomes (B).

### While Ineffectual in Initial PRC2 Recruitment, EPOP Facilitates H3K27me3 Deposition de novo

We found that EPOP stimulated PRC2 HMT activity *in vitro* (**Figure 1**), yet previous work suggested that EPOP inhibits PRC2-mediated deposition of H3K27me3 on chromatin. Notably, these latter experiments involved the knockdown of EPOP in cells under “steady-state” conditions wherein H3K27me3-domains were already established^13,14^ and therefore, did not probe for EPOP-mediated effects on the *de novo* establishment of PRC2-repressive domains. Our HMT experiments suggested that, in the absence of any confounding factors, such as other cofactors and pre-established H3K27me3, EPOP stimulated PRC2 HMT activity (**Figure 1**). To corroborate these findings within a cellular context, we employed the previously developed “EED rescue system”^7^. This system entails the full depletion of PRC2 and its associated H3K27me3-domains through the knockout (KO) of the core-PRC2 subunit, EED, in mESCs. PRC2 is then restored through EED re-expression and the kinetics of H3K27me3 deposition *de novo* are monitored using ChIP-seq.

To probe for EPOP-mediated effects on the *de novo* recruitment of PRC2 to chromatin, we used CRISPR-Cas9 to generate an EPOP KO mESC line similar to the MTF2 KO, JARID2 KO, and MTF2/JARID2 double KO (dKO) lines comprising the EED-rescue system previously established^7^. We then performed ChIP-seq for HA-EED at 0 h and 24 h after EED rescue to directly monitor the initial recruitment of PRC2 to chromatin as a function of the loss of EPOP, MTF2 and/or JARID2. Consistent with prior findings, MTF2 loss severely impaired HA-EED recruitment, JARID2 loss had only a modest effect, and the combined knockout nearly abolished recruitment^7^. By contrast, EPOP KO cells exhibited PRC2 binding comparable to wild type, demonstrating that EPOP neither promotes nor inhibits the initial recruitment of PRC2 (**Figure 2A**), despite its strong enrichment at nucleation sites^7^.

**Figure 2.**
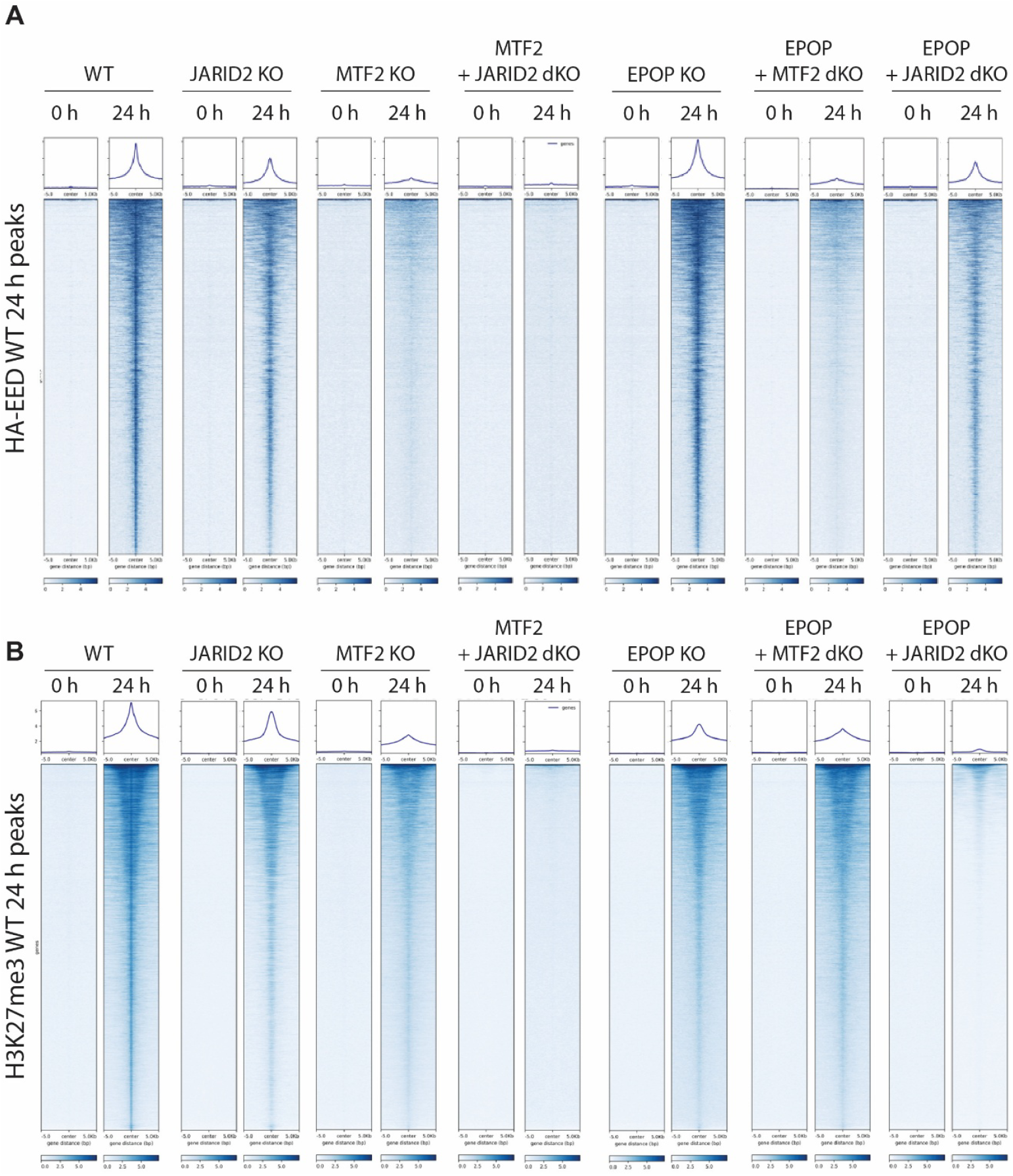
EPOP facilitates H3K27me3 deposition *de novo* while not required for initial PRC2 recruitment. (A and B) Heat maps of ChIP-seq peaks for HA-EED (A) and H3K27me3 (B) within a 20 kb window centered on the maximum value of peak signal in mESCs with the genotypes of cells engineered with the EED rescue system as indicated. Cells were collected at 0 h and 24 h after EED re-expression.

We speculated that loss of EPOP might unmask a role in PRC2 recruitment when combined with the loss of other cofactors, as seen with JARID2 and MTF2^7^. To test this, we generated EPOP/MTF2 and EPOP/JARID2 dKO cell lines in the EED-rescue background and performed HA-EED ChIP-seq. EPOP/MTF2 dKO cells recapitulated the severe loss of recruitment observed in cells with MTF2 KO alone, and EPOP/JARID2 dKO cells showed the same modest reduction as in JARID2 KO cells, affirming that EPOP has no measurable impact on *de novo* PRC2 recruitment (**Figure 2A**). These results established that EPOP is dispensable for *de novo* PRC2 recruitment.

Importantly, our result differs from prior studies in which knockdown (KD) of EPOP led to an increase in PRC2 chromatin occupancy and elevated H3K27me3 at EPOP-bound sites, leading to the interpretation that EPOP restrains PRC2 binding^13,14^. Notably, those experiments were performed under steady state conditions wherein PRC2 domains and H3K27me3 landscapes were already established in mESCs. Under these conditions, loss of EPOP allowed for an increased occupancy of the PRC2 core subunit, SUZ12, and H3K27me3 catalysis, coincident with enhanced binding of JARID2 at the same sites^13^. As EPOP and JARID2 define biochemically distinct PRC2 subcomplexes^13,20^, this shift likely reflects the functional compensation of PRC2-EPOP by PRC2–JARID2. Given the more prominent role of JARID2 in promoting PRC2 recruitment compared to EPOP as demonstrated here (**Figure 2A**), this switch provides a plausible explanation for the increased PRC2 occupancy observed upon EPOP depletion in steady-state cells. In contrast, our EED-rescue system interrogates *de novo* recruitment in the absence of any pre-existing PRC2 domains and under these circumstances, we find that EPOP neither promotes nor inhibits the initial chromatin binding of PRC2.

Although EPOP is dispensable for the initial recruitment of PRC2 in cells, it does indeed have a robust stimulatory effect on PRC2 HMT activity *in vitro*. We therefore speculated that EPOP might also contribute to the *de novo* deposition of H3K27me3 in a cellular context. To this end, we performed ChIP-seq for H3K27me3 in the EED-rescue system using the same panel of KO cell lines. Consistent with our previous findings^7^, MTF2 KO led to a strong reduction in H3K27me3 with residual signal that was almost completely lost in the MTF2/JARID2 dKO, whereas JARID2 KO alone had only a mild effect (**Figure 2B**). Strikingly, EPOP KO led to a substantial reduction in H3K27me3 deposition (∼50% decreased relative to WT), though less severe than that observed in the case of MTF2 KO. Importantly, EPOP/MTF2 dKO cells resembled the MTF2 KO alone, while EPOP/JARID2 dKO cells showed a severe loss of H3K27me3, almost similar to the case of MTF2/JARID2 dKO (**Figure 2B**). Thus, EPOP, in conjunction with MTF2 and JARID2, promotes the *de novo* deposition of H3K27me3 *in vivo*.

### EPOP and MTF2 Regulate PRC2 HMT Activity in a DNA-Sequence-Specific Manner

Building on our finding that EPOP contributes to *de novo* H3K27me3 deposition *in vivo* (**Figure 2**), we developed a defined *in vitro* system to gain mechanistic insight into this process. In this case, nucleosomes are assembled free of pre-existing modifications, thereby modeling the process of *de novo* H3K27 methylation. Notably, CpG islands at PRC2 nucleation sites are highly enriched for tandem GA- and GCN-repeat motifs compared to other genomic regions^7^. These motifs resemble binding sequences recognized by PRC2 cofactors, such as JARID2 and MTF2, respectively^8,9^, suggesting that they might influence subcomplex-specific regulation of PRC2 activity. To test this possibility, we reconstituted recombinant dinucleosomes containing two 601-positioning sequences separated by a 40 bp linker, a configuration previously shown to maximally stimulate PRC2 HMT activity^21^. Into the linker region, we inserted either a random sequence (R), a GA repeat, or a GCN repeat, thereby generating substrates that mimic chromatin features of PRC2-specific nucleation sites. Using these R-, GA-, and GCN-dinucleosomes, we assessed how PRC2-EPOP and PRC2-MTF2 respond to the context of DNA-sequence in promoting H3K27 methylation.

When incubated with dinucleosomes containing random linker DNA (R-dinucleosomes), PRC2-MTF2 showed a substantial increase in HMT activity relative to PRC2-core. Strikingly, PRC2-EPOP also showed a substantial increase in HMT activity that was approximately equal to that of MTF2 (**Figures 3A-B**). Thus, consistent with our findings above (**Figure 1**), EPOP robustly stimulated the HMT activity of PRC2 on dinucleosomes. With GCN-dinucleosomes, the HMT activity of the PRC2-core and of PRC2-EPOP was similar to that with R-dinucleosomes. However, PRC2-MTF2 HMT activity was stimulated approximately two-fold compared to its activity on R-dinucleosomes (**Figures 3C-D**). Previous reports have demonstrated that the winged-helix domain of MTF2 and other PCL proteins enhance their affinity for GC-rich DNA^9^. In accordance, our data showed that such enhancement is demonstrable in the case of nucleosomes bridged by GCN-rich DNA, which specifically and robustly enhanced the HMT activity of PRC2-MTF2.

**Figure 3.**
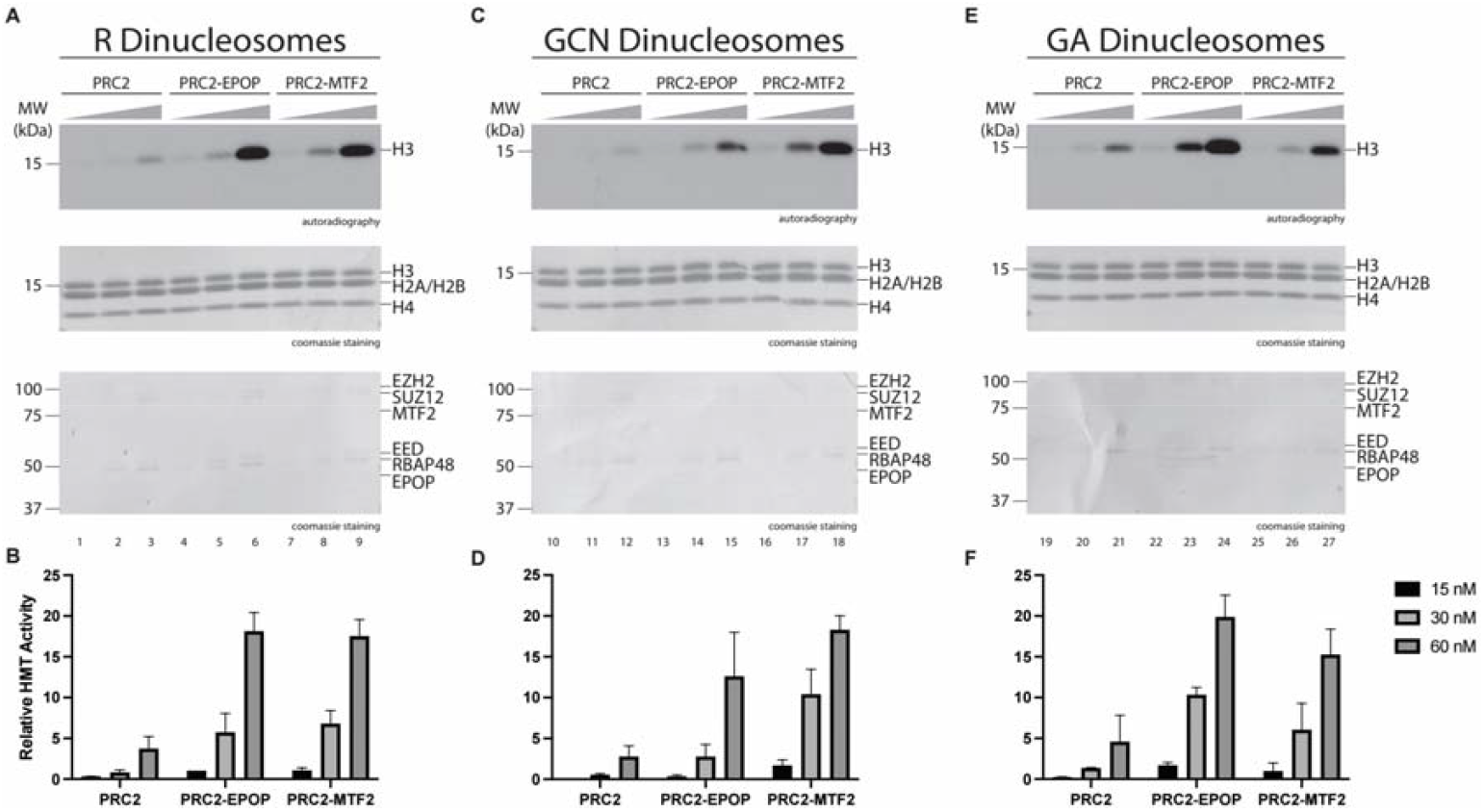
Regulation of PRC2 Histone Methyltransferase Activity by MTF2 and EPOP is DNA Sequence-specific. (A, C, E) HMT assays (30 min incubation) performed with increasing amounts of PRC2, PRC2-EPOP, and PRC2-MTF2 (15, 30, or 60 nM) in the context of dinucleosomes (300 nM) containing distinct versions of a 40 bp linker DNA sequence: (A) R (random), (C) GCN tandem repeats, or (E) GA tandem repeats. Top image: H3 methylation levels shown by autoradiography. Middle and bottom images: histones from dinucleosomes and subunits of PRC2 complexes shown by Coomassie blue staining of SDS-PAGE gels, respectively. (B, D, and F) Quantification of relative amounts of ^3^H-SAM incorporated into H3 from HMT assays performed in A, C, and E, respectively, normalized to PRC2-EPOP (15 nM) on R dinucleosomes.

Conversely, with GA-rich dinucleosomes, the HMT activities of PRC2-MTF2 and PRC2-core were similar to their respective activity with R-dinucleosomes. However, PRC2-EPOP HMT activity was stimulated approximately two-fold compared to its activity on R-dinucleosomes (**Figures 3E-F**), substantiating our findings above that EPOP exhibited a robust stimulatory effect on PRC2 HMT activity, consistent with its enrichment at PRC2 nucleation sites^7^. Interestingly, EPOP is thought to be largely unstructured and devoid of DNA-binding motifs^14^, such that its preference for GA-rich linker DNA was unexpected. Nonetheless, EPOP recognized GA-dinucleosomes and robustly enhanced PRC2 HMT activity through a yet to be determined mechanism. Notably and to the best of our knowledge, EPOP is the first cofactor shown to confer sequence specificity by enhancing PRC2 HMT activity on GA-rich dinucleosomes, while broadly stimulating activity across substrates, suggesting a role in promoting *de novo* H3K27me3 deposition at GA motif–containing nucleation sites during development (see Discussion). Thus, EPOP and MTF2 moderate PRC2 HMT activity in a DNA-sequence specific manner.

### EPOP and MTF2 Differentially Enhance PRC2 Affinity in a Sequence- and Chromatin-Dependent Manner

We next sought to determine the mechanistic basis by which EPOP and MTF2 stimulate PRC2 HMT activity. Given that other cofactors, such as JARID2 and AEBP2^15,21^, enhance PRC2 HMT activity by increasing PRC2 binding affinity for nucleosomes, we speculated that EPOP and MTF2 may operate similarly. Thus, we performed electrophoretic mobility shift assays (EMSA) using the same reconstituted dinucleosome substrates as in Figure 3. With R-dinucleosomes, both EPOP and MTF2 strongly increased the binding affinity of PRC2 toward nucleosomes relative to PRC2-core (**Figures 4A-B and S1A**). Therefore, MTF2 and EPOP likely increased PRC2 HMT activity by enhancing its affinity toward nucleosomes. On GA-linker dinucleosomes, PRC2–EPOP bound with markedly higher affinity than PRC2–MTF2 or core (**Figures 4C-D and S1B**), reflecting its enhanced HMT activity on this substrate (**Figure 3**). Notably, PRC2-EPOP maintained strong binding on GA-dinucleosomes relative to R-dinucleosomes, whereas PRC2–MTF2 binding was attenuated on GA-dinucleosomes compared with R-dinucleosomes. Together, the superior binding and catalytic activity of PRC2–EPOP on GA-dinucleosomes suggest that EPOP preferentially facilitates H3K27me3 deposition *de novo* at GA⍰ enriched nucleation sites. On GCN-linker dinucleosomes, PRC2–MTF2 exhibited enhanced binding relative to R-dinucleosomes (**Figures 4E-F and S1C**), contrasting with its reduced affinity on GA-dinucleosomes. By comparison, PRC2–EPOP maintained consistently strong binding across R, GA, and GCN substrates, while PRC2-core remained weakest. The modestly stronger binding of PRC2–MTF2 relative to PRC2-EPOP on GCN-dinucleosome aligns with its highest HMT activity on this substrate (**Figure 3**), and suggests that MTF2 may contribute to H3K27me3 deposition *de novo* at nucleation sites enriched for GCN motifs.

**Figure 4.**
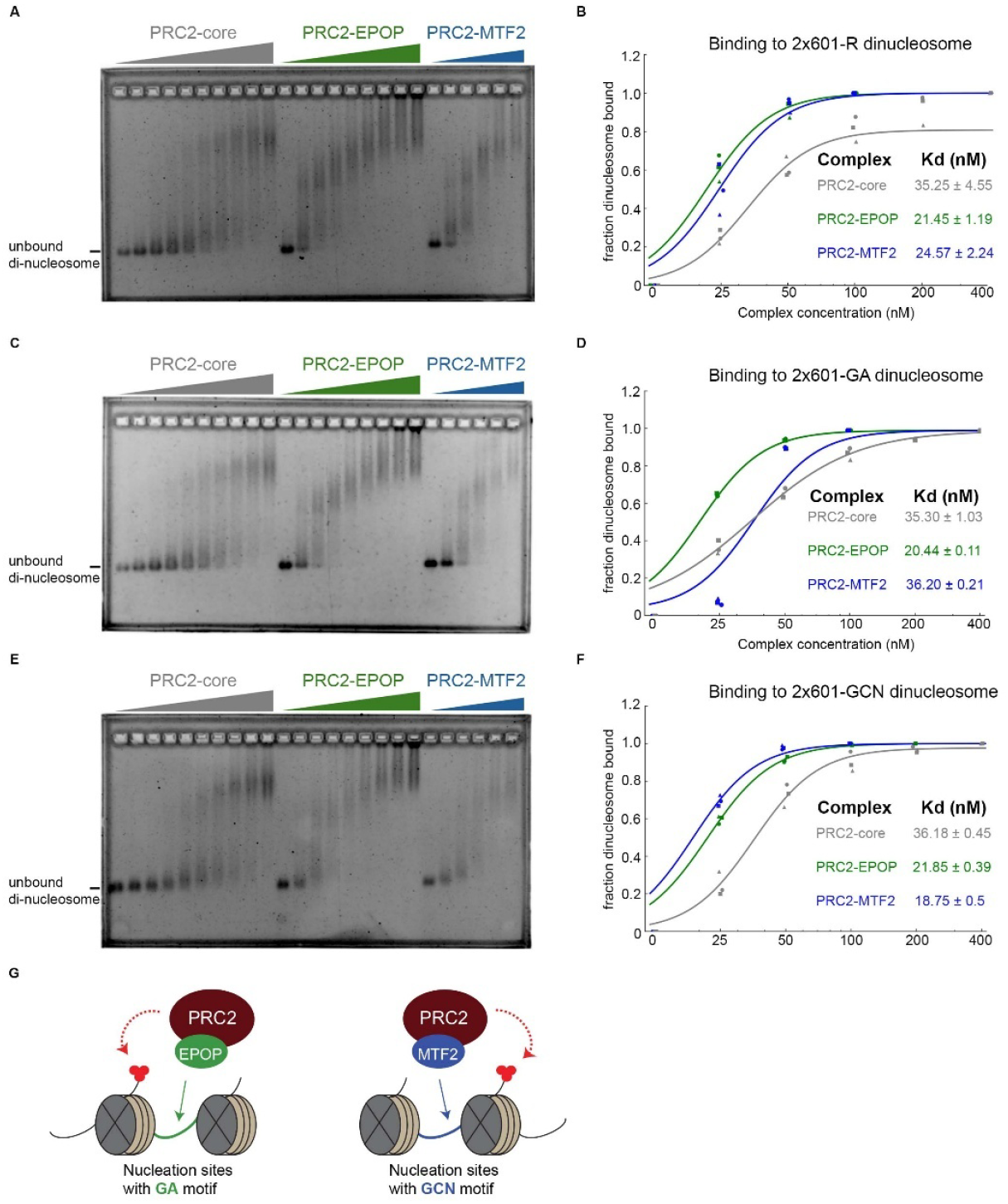
EPOP and MTF2 Promote PRC2 Binding to Dinucleosomes, with Preferential Affinity of PRC2–EPOP for GA Linkers. (A, C, E) EMSAs examining binding between PRC2-core, PRC2–EPOP, or PRC2–MTF2 and dinucleosomes reconstituted with two Widom 601 positioning sequences separated by a 40 bp linker. Linker identities: random sequence (R; A), GA tandem repeats (C), or GCN tandem repeats (E). Reactions were incubated for 30 min and resolved on agarose gels; DNA was visualized by SYBR Gold staining. (B, D, F) Quantitative binding analysis for R-(B), GA-(D), and GCN-(F) linker dinucleosomes, measured from the other half of each reaction performed in A, C, and E, respectively, resolved on acrylamide gels for increased sensitivity. The corresponding acrylamide gels are shown in Figure S1A–C. Data were fit with a sigmoidal binding function to calculate dissociation constants (Kd). *n* = 3 independent experiments. (G) Schematic diagram depicting EPOP- and MTF2-mediated *de novo* H3K27me3 deposition at GA- and GCN-enriched nucleation sites, respectively.

To test whether the sequence preferences of PRC2–EPOP and PRC2–MTF2 arise from the linker DNA itself or require a chromatin context, we performed EMSAs using the DNA fragments employed for dinucleosome reconstitution (R, GA, and GCN). Both EPOP and MTF2 enhanced PRC2 binding to DNA relative to PRC2-core, without exhibiting sequence specificity (**Figures S2-3**). Across all three DNA substrates, PRC2–MTF2 consistently bound more strongly than PRC2–EPOP, which in turn exceeded PRC2-core. These findings demonstrate that the distinct GA and GCN preferences observed with dinucleosomes (**Figure 4**) are not solely due to accessory factor recognition of specific DNA sequences, but instead emerge within a chromatin context, consistent with the physiological setting of PRC2 nucleation sites.

## Discussion

Much effort has been devoted towards developing a framework to account for the genome-wide binding patterns of PRC2 and the concomitant presence of its product, H3K27me3, in mammalian cells. Recent work provided insight into this process by demonstrating that PRC2 is recruited *de novo* to a discrete set of nucleation sites to initiate and then spread H3K27me3^7^. However, the relationship between the GCN- and GA-tandem repeats enriched at these sites and the cofactors that guide PRC2 to these foci has remained unclear. Although previous studies have consistently demonstrated the central role of MTF2 and other cofactors in promoting recruitment of PRC2 and deposition of H3K27me3^9,15–19^, a consensus regarding the function of EPOP in coordinating these activities has not been reached. That EPOP is highly enriched at PRC2 nucleation sites^7^ would suggest that it facilitates PRC2 recruitment and its subsequent catalysis of H3K27me3. Yet, previous studies suggested that EPOP inhibits the recruitment of PRC2 and its deposition of H3K27me3^13,14^. However, these studies focused on the effect of knocking-down EPOP in “steady-state” cells wherein H3K27me3-domains have already been established, rather than the role of EPOP in coordinating the *de novo* recruitment and deposition of H3K27me3. Thus, these previous findings may instead relate to the function of EPOP in stabilizing PRC2 and H3K27me3 on chromatin. Indeed, alternative explanations can account for the observations derived from these experiments. Although MTF2 and EPOP associate with the PRC2-core complex through non-overlapping binding sites on its SUZ12 subunit^22,23^, the absence of EPOP would likely foster the further association of the PRC2-core complex with its remaining cofactors, such as MTF2 and JARID2. This scenario would likely promote the efficient targeting of PRC2 to chromatin^22,23^. Indeed, given that MTF2 is the strongest driver of PRC2 recruitment^7,10,11^, this likelihood could result in higher levels of chromatin-bound PRC2 and higher levels of H3K27me3 in the absence of EPOP, thereby apparently disqualifying EPOP as a positive regulator of PRC2 recruitment and activity. Instead, our findings here show that EPOP neither inhibits nor promotes the initial recruitment of PRC2 to chromatin.

Previous experiments have interpreted MTF2 as being the only PRC2 cofactor whose absence in mESCs results in a significant disruption to PRC2 recruitment, as the absence of other cofactors, such as JARID2 and AEBP2, presents no appreciable defect^7,10,11^. These findings are consistent with those in our current study, in which we find PRC2 recruitment to be unperturbed in EPOP KO cells (**Figure 2A**). However, given that EPOP robustly stimulates PRC2 HMT activity (**Figures 1** and **3**), these findings do not preclude the possibility that EPOP stimulates the activity of PRC2 on chromatin in a cellular context. For example, it is well-established that AEBP2 and JARID2 have robust stimulatory effects on PRC2^15–19,21^ and are essential for proper patterning of H3K27me3 on chromatin, yet lack an effect in coordinating PRC2 recruitment, relative to MTF2^7^. In this study, we provide evidence that EPOP does aid PRC2 in H3K27me3 deposition *de novo* on chromatin (**Figure 2B**).

While PRC2-EPOP is stimulated to the greatest extent in the presence of GA-dinucleosomes, PRC2-MTF2 is stimulated most highly in the presence of GCN-dinucleosomes, as shown here (**Figure 3**). These data reveal a regulatory paradigm whereby PRC2 accessory subunits stimulate the HMT activity of PRC2 in a DNA-sequence specific manner. It will be important to assess if GCN- and GA-sequences differentially recruit MTF2 and EPOP, respectively, in cells. Also, it will be informative to evaluate the effect that the individual knock-out of EPOP and MTF2 has on the *de novo* deposition of H3K27me3 at GA- and GCN-rich nucleation sites, in particular.

The data presented here provide important insights into the mechanistic basis by which PRC2 executes unique patterns of H3K27me3 across developmental lineages^15–17,24–26^, given that PRC2 cofactor expression patterns are cell type-specific^27^. For example, cells expressing EPOP but not MTF2 might robustly catalyze H3K27me3 at sites enriched with GA-repeats and consequently maintain transcriptional repression within these domains. However, in the absence of MTF2, H3K27me3 would be diluted at GCN-rich sites resulting in transcriptional derepression of the associated genes. We expect these regulatory processes to be executed early in development, as the expression of many PRC2 accessory subunits diminishes at later developmental stages^15–17,24–26^. Future experiments should therefore probe the functional role of EPOP, MTF2, and other PRC2 cofactors in maintaining H3K27me3 patterns in differentiated cellular states.

This study also contributes mechanistic insights as to how accessory factors stimulate the HMT activity of PRC2. Most cofactors, like AEBP2 and JARID2, enhance binding of PRC2 to nucleosomes^15,21^, which is believed to elicit a stimulatory effect on PRC2 HMT activity. Similarly, we showed that MTF2 stimulates PRC2 HMT activity, likely by increasing the binding affinity of PRC2 toward nucleosomes. Moreover, the binding affinity of PRC2-MTF2 is further enhanced by GCN-dinucleosomes (**Figure 4**), which likely explains the robust stimulatory effect on PRC2-MTF2 HMT activity with this substrate (**Figure 3**). Surprisingly, while EPOP also likely stimulates PRC2 HMT activity by increasing the binding affinity of PRC2 toward nucleosomes (**Figure 4**), EPOP is believed to be a largely unstructured protein devoid of any known DNA binding domains^14^. The mechanistic basis by which GA-rich nucleosomal DNA provides additional stimulation of PRC2 HMT activity in this case (**Figure 3**) remains unclear, as the binding affinity of PRC2-EPOP appears similar for GA-dinucleosomes and control R-dinucleosomes (**Figure 4**). However, it is still noteworthy that PRC2-MTF2 shows decreased binding relative to PRC2-EPOP on GA-dinucleosomes, which, in a cellular context, may have mechanistic implications in determining how these two PRC2 subcomplexes differentially recognize unique sequence motifs at nucleation sites. Notably, supplemental EMSAs with the corresponding naked DNA templates revealed that both EPOP and MTF2 enhanced PRC2–DNA binding without exhibiting sequence specificity (**Figure S2**), underscoring that the observed GA and GCN preferences arise specifically in the chromatin context of nucleosomes. Of note, EPOP-induced stimulation of PRC2 HMT activity may occur through alternative routes, as accessory factors may also participate in the allosteric activation of PRC2. For example, it was shown that similar to H3K27, JARID2-K116 is methylated (JARID2-K116me3) by the catalytic subunit of PRC2, EZH2, and similar to H3K27me3, JARID2-K116me3 allosterically stimulates the HMT activity of PRC2 through association with its EED subunit^18^. Furthermore, it was recently shown that in addition to enhancing PRC2-nucleosome binding, PALI1 is also methylated by PRC2 and allosterically stimulates its HMT activity^28^. Thus, the possibility exists that EPOP is methylated by EZH2, which can then be recognized by EED with a resultant allosteric activation of the complex. Nonetheless, the findings presented here support a framework whereby cell-type specific patterns of H3K27me3 can arise from lineage-specific cofactor expression patterns^27^. Future investigations should address this possibility to help clarify the key processes by which PRC2 initiates and maintains unique patterns of H3K27me3 throughout development.

## Materials and Methods

### Cell lines and culture condition

All ESC lines (E14 and derivatives) were grown in DMEM supplemented with 15% FBS, L-glutamine, penicillin/streptomycin, sodium pyruvate, non-essential amino acids, 0.1 mM β-mercaptoethanol, LIF, and 2i inhibitors, which include 1 μM MEK1/2 inhibitor (PD0325901) and 3 μM GSK3 inhibitor (CHIR99021) on 0.1% gelatin coated plates.

### Purification of PRC2 subcomplexes using Baculovirus expression system

To purify human PRC2 subomplexes containing EPOP and MTF2, the biGBac cloning system was first used assemble all subunits into a single plasmid (1). A plasmid encoding all four of the core PRC2 subunits (3x-Strep-EZH2, FLAG-SUZ12, 6x-His-EED, 6x-His-RBAP48) was previously constructed through biGBac cloning by the laboratory of Karim-Jean Armache and was generously shared with us. We then used biGBac cloning to combine the core subunits with Strep-MTF2 and 6x-His-EPOP to assemble two additional plasmids encoding PRC2-MTF2 and PRC2-EPOP. Standard procedures were then used to generate bacmids and baculoviruses, such that single baculoviruses encoded all subunits of each PRC2 subcomplex. 500 mL of Sf9 cells were infected with baculoviruses at a 1:100 ratio (5 mL) and incubated for 60 h. The cells were then harvested and resuspended in BC150 buffer (25 mM Hepes-NaOH, pH 7.5, 1 mM EDTA, 300 mM NaCl, 5% glycerol, 0.2 mM DTT, and 0.1% NP-40) with protease inhibitors (1 mM phenylmethlysulfonyl fluoride (PMSF), 0.1 mM benzamidine, 1.25 mg/ml leupeptin and 0.625 mg/ml pepstatin A) and phosphatase inhibitors (20 mM NaF and 1 mM Na3VO4). Cells were then lysed by sonication (Fisher Sonic Dismembrator model 100). PRC2 subcomplexes were then purified through Ni-NTA agarose resin (Qiagen) (for PRC2-core and PRC2-EPOP) or Strep-Tactin Macroprep resin (IBA/Neuromics) (for PRC2-MTF2) and dialyzed against BC150. His-TEV and GST-HRV-3C proteases were subsequently used to cleave the affinity tags from each subunit. The samples were then incubated with Glutathione Sepharose 4 Fast Flow resin (MilliporeSigma) to remove GST-HRV-3C. The flowthrough was then subjected to size exclusion chromatography (Superose 6 XK 16-70 (MilliporeSigma)) to remove contaminants, His-TEV, and any remaining GST-HRV-3C. The peak fractions were then pooled and concentrated.

### Nucleosome reconstitution

Recombinant histones were generated as previously described. Briefly, each core histone was expressed in Rosetta (DE3) cells (Novagen), extracted from inclusion bodies, and purified by sequential anion and cation chromatography. For refolding recombinant octamers, equal amounts of histones were mixed and dialyzed into refolding buffer (10 mM Tris-HCl, pH 7.5, 2 M NaCl, 1 mM EDTA, and 5 mM β-mercaptoethanol). Octamers were further purified by size exclusion chromatography on a 24-ml Superdex 200 column (GE healthcare) in refolding buffer. Recombinant oligonucleosomes were reconstituted by sequential salt dialysis of octamers and plasmid having 12 repeats of the 601-nucleosome positioning sequence. Recombinant dinculeosomes were assembled in a similar fashion using a DNA template consisting of two 601-sequences separated by 40 base pairs of DNA consisting of a random control sequence, GCN-tandem repeats, or GA-tandem repeats.

For salt dialysis, DNA was mixed with titrated histone octamers and assembled using dialysis from High salt buffer (2 M NaCl, 10 mM Tris-HCl pH 7.5, 1 mM EDTA, 1 mM 2-Mercaptoethanol) to No salt buffer (10 mM Tris-HCl pH 7.5, 1 mM EDTA, 1 mM 2-Mercaptoethanol). The reactions were dialyzed gradually from High salt buffer (1 liter) to No salt buffer (150 ml) at a flow rate of 1 ml/min using a peristaltic pump. After dialysis for about 16 h, buffer was exchanged into No salt buffer (10 mM Tris-HCl pH 7.5, 1 mM EDTA, 1 mM 2-Mercaptoethanol) and allowed to dialyze for 3 h. Nucleosomes were then transferred to LoBind tubes and stored at 4 °C. After reconstitution, the samples were analyzed using 5% native acrylamide gel or a 1.2% agarose gel stained by SYBR Gold.

### HMT assay

Standard HMT assays were performed in a total volume of 15 μl containing HMT buffer (50 mM Tris-HCl, pH 8.5, 5 mM MgCl2, and 4 mM DTT) with the indicated concentration of 3H-labeled S-Adenosylmethionine (SAM, Perkin Elmer), 300 nM of recombinant oligonucleosomes, and recombinant human PRC2 complexes. The reaction mixture was incubated for 60 min (for experiments in Chapter 2) or 30 min (for experiments in Chapter 3) at 30 °C and stopped by the addition of 4 μl SDS buffer (0.2 M Tris-HCl, pH 6.8, 20% glycerol, 10% SDS, 10 mM β-mercaptoethanol, and 0.05% Bromophenol blue). For Chapter 2, a titration of PRC2 (from 5 to 60 nM) was performed under these conditions to optimize the HMT reaction within a linear range, and the yield of each HMT reaction was measured using the following procedures. This was also done for the work in Chapter 3, except that PRC2 subcomplexes were titrated at concentrations of 15 nM, 30 nM, and 60 nM. After HMT reactions, samples were incubated for 5 min at 95 °C and separated on SDS-PAGE gels. The gels were then subjected to Coomassie blue staining for protein visualization or wet transfer of proteins to 0.45 μm PVDF membranes (Millipore). The radioactive signals were detected by exposure on autoradiography films (Denville Scientific).

### Electrophoretic mobility shift assay (EMSA)

Reconstituted nucleosomes or free DNA were mixed with PRC2 subcomplexes at desired concentration using 5xEMSA buffer (100 mM Tris-HCl pH 7.5, 500 mM NaCl, 12.5 mM MgCl2, 0.5 mM ZnCl2, 0.5mg/ml BSA, 10 mM 2-mercaptoethanol, 0.25% NP-40, 25% Glycerol).Binding was carried out at 30 °C for 30 min, then subjected to non-denaturing gel electrophoresis at 6.6□V/cm over a 1.2% agarose gel buffered with 1× TBE at 4□°C for 1.5□h or at 100□V over a fresh prepared 5% native acrylamide gel buffered with 0.25× TBE at 4□°C for 2.5□h. Gels were stained with SYBR Gold at RT and then imaged. The fractions of bound nucleosomes or DNA were calculated based on the unbound nucleosomes or DNA band with the densitometry analysis carried out using ImageJ. Because of the higher sensitivity in 5% native acrylamide gel compared to agarose gel, data from the native acrylamide gel were fitted into sigmoidal curve with Hill slope function in RStudio. All experiments were performed in triplicate.

### CRISPR-mediated genome editing

To generate stable *Epop* KO cell lines from either WT, *Jarid2* KO or *Mtf2* KO, sgRNAs were designed using CRISPR design tool in https://benchling.com. sgRNAs in Table S1 were cloned in pSpCas9(BB)-2A-GFP (PX458, a gift from Feng Zhang, Addgene plasmid #48138) and transfected into mESCs, using Lipofectamine 2000 (Life Technologies). GFP-positive cells were sorted 48 hr after transfection and 20,000 cells were plated on a 15 cm dish. Single mESC was allowed to grow to a colony for ∼5 days and then was picked, trypsinized in Accutase for 5 min, and split into two individual wells of two 96-well plates for genotyping and culture, respectively. Genomic DNA was extracted using lysis buffer (50 mM Tris-HCl, pH 8, 2 mM NaCl, 10 mM EDTA, 0.1% SDS) supplemented with protease K. The resulting PCR products were sent for sequencing to determine the presence of a deletion or a mutation event. Clones were further confirmed by western blot.

### ChIP-seq library preparation

For cross-linking, ESCs were fixed in 1% formaldehyde for 10 min at RT directly on plates and quenched with 125 mM glycine for 5 min at RT.

Cell pellets were washed twice in PBS and nuclei were isolated using buffers in the following order: LB1 (50 mM HEPES, pH 7.5 at 4°C, 140 mM NaCl, 1 mM EDTA, 10% Glycerol, 0.5% NP40, 0.25% Triton X; 10 min at 4°C), LB2 (10 mM Tris, pH 8 at 4°C, 200 mM NaCl, 1 mM EDTA, 0.5 mM EGTA; 10 min at 4°C), and LB3 (10 mM Tris, pH 7.5 at 4°C, 1 mM EDTA, 0.5 mM EGTA, and 0.5% N-Lauroylsarcosine sodium salt). Chromatin was fragmented to an average size of 250 bp in LB3 buffer using a Diagenode Bioruptor. 200 μg sonicated chromatin, 4 ug antibody and 20 ul Dynabeads were used in each ChIP reaction supplemented with 0.5x volumn of incubation buffer (3% Triton X, 0.3% Na Deoxycholate, 15 mM EDTA). 1 μg of Drosophila chromatin and 0.2 μg of anti-Drosophila H2A.X antibody were added in each ChIP reaction as spike-in references. After 5 consecutive washes with RIPA buffer (50 mM HEPES, pH 7.5 at 4°C, 0.7% Na Deoxycholate, 1 mM EDTA, 1% NP40, 500 mM LiCl) and one wash with TE+50 mM NaCl, the beads-bound DNA was eluted in freshly prepared elution buffer (50 mM Tris, pH 8, 10 mM EDTA, 1% SDS) at 65°C for 20 min. Eluted DNA was de-crosslinked at 65°C overnight, followed by protease K and RNase A treatment.

For Library preparation, IP’ed DNA (∼1-30 ng) was end-repaired using End-It Repair Kit, tailed with deoxyadenine using Klenow exo-, and ligated to custom adapters with T4 Rapid DNA Ligase (Enzymatics). Fragments of 200-600 bp were size-selected using Agencourt AMPure XP beads (0.5X and 0.3X), and subjected to PCR amplification using Q5 DNA polymerase. Libraries were size-selected using Agencourt AMPure XP beads (0.75X), quantified by Qubit™ dsDNA HS Assay Kit and quality checked by High Sensitivity D1000 ScreenTape. Libraries were sequenced on the Illumina NovaSeq 6000 platform.

## QUANTIFICATION AND STATISTICAL ANALYSIS

### ChIP-seq data analysis

ChIP-seq data analysis was performed as described previously^35^. Briefly, reads were aligned to the mouse reference genome mm10 and dm6 for spike-in samples, using Bowtie2 with default parameters. Reads of quality score less than 30 were removed using samtools and PCR duplicates were removed using picard. Regions in mm10 genome blacklist was removed using bedtools and bigwig files were generated using deeptools and parameters: --binSize 50 --normalizeUsing RPKM --ignoreDuplicates -- ignoreForNormalization chrX --extendReads 250 for visualization in IGV. Peaks were called using MACS3 with parameters: -g mm --keep-dup 1 --nomodel --extsize 300. For visualization of ChIP-seq, uniquely aligned reads mapping to the mouse genome were normalized using dm6 spike-in as described previously^36^. Heatmaps were performed using the functions computeMatrix followed by plotHeatmap and plotProfile from deepTools.

## DATA AND CODE AVAILABILITY

The accession numbers for the raw data FASTQ files and processed files for all sequencing data are deposited in NCBI GEO are GEO: GSE306783. Reviewers can access the data at https://www.ncbi.nlm.nih.gov/geo/query/acc.cgi?acc=GSE306783 using token “ebsrweuwrhsbvsd”.

## Acknowledgments

We thank Drs. L. Vales for critical reading and editing of the manuscript, H. Bharti for technical assistance with nucleosome reconstitution, as well as past and current Reinberg laboratory members for critical comments and discussions. We also thank the K.J. Armache laboratory for providing the biGBac plasmid and guidance on cloning, the New York University Langone Medical Center (NYULMC) Genome Technology Center for help with sequencing, the NYULMC Cytometry and Cell Sorting Core for help with FACS. This study utilized computing resources at the High-Performance Computing Facility of the Center for Health Informatics and Bioinformatics at the NYULMC. This work was supported by grants to D.R. from the NIH (R01NS100897, R01CA199652) and the Howard Hughes Medical Institute.

## Author Contributions

J.G. conceived the study, designed and developed the core experimental assays, performed the majority of the experiments, and drafted the initial version of the manuscript; S.L. performed the EMSA assay, conducted the bioinformatic analyses, and led the revision of the manuscript during the submission and review process; L.P. helped with nucleosome reconstitution and EMSA assay; O.O. helped with generating knockout cell lines in EED rescue system; D.R. supervised the project and provided strategic guidance throughout. All authors contributed to data interpretation and manuscript editing, and approved the final version of the manuscript.

## Competing Interest Statement

The authors declare no competing interests.

## Figures and Tables

**Figure S1.**
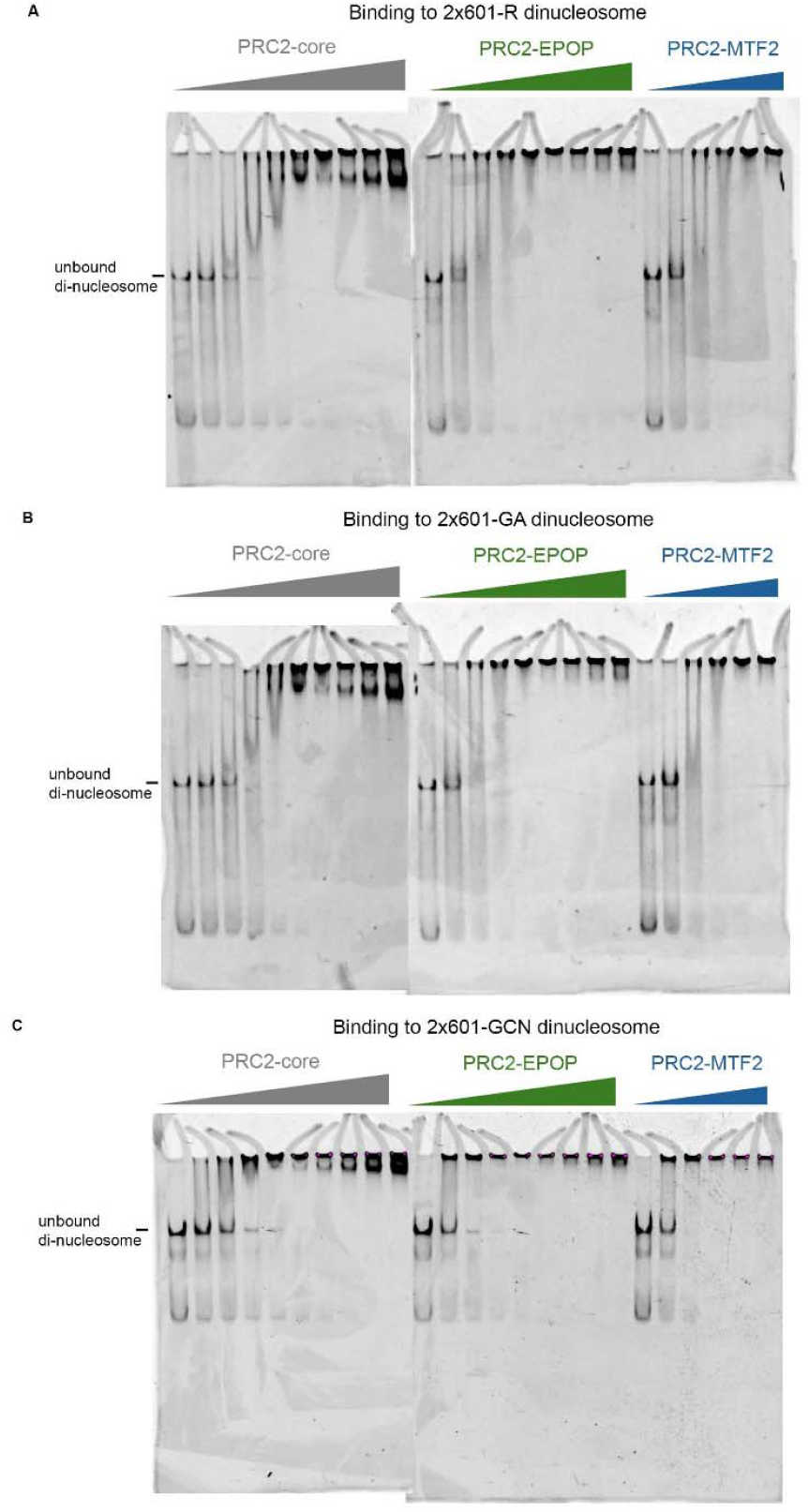
EMSAs resolved on acrylamide gels for PRC2–core, PRC2–EPOP, and PRC2–MTF2 binding to dinucleosomes with defined linker sequences, corresponding to Fig. 4. (A–C) EMSAs of PRC2–core, PRC2–EPOP, and PRC2–MTF2 incubated with dinucleosomes containing two Widom 601 positioning sequences separated by a 40 bp linker comprising either a random sequence [R; (A)], GA tandem repeats (B), or GCN tandem repeats (C). For each condition, half of the reactions shown in Figure 4A, C, and E were resolved on agarose gels (Figure 4), while the other half were resolved in parallel on acrylamide gels to provide increased detection sensitivity. DNA was visualized by SYBR Gold staining. These acrylamide gels were used for quantitative analysis and curve fitting shown in Fig. 4B, D, and F.

**Figure S2.**
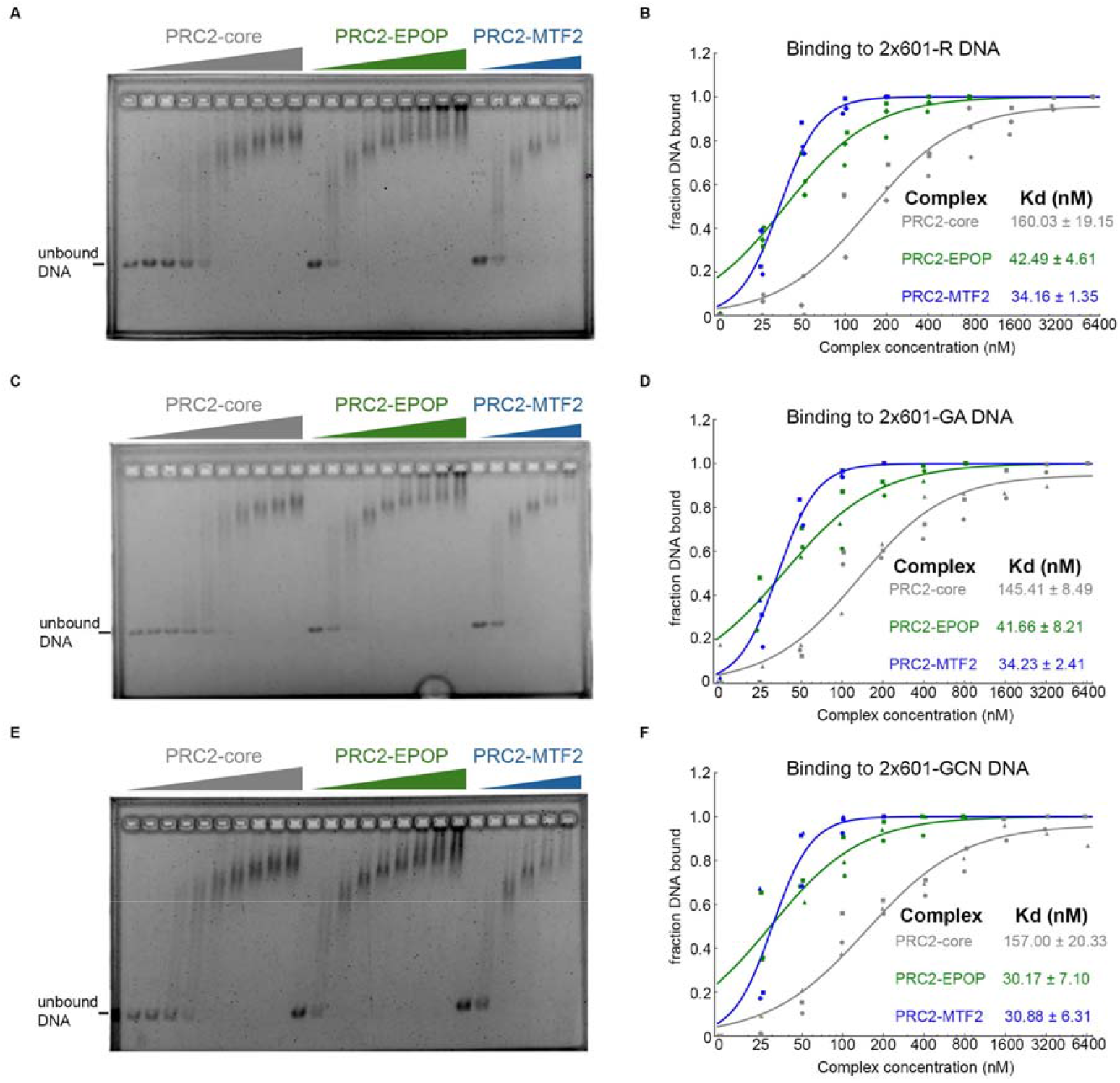
EPOP and MTF2 Enhance PRC2 Binding to Free DNA Independent of Linker Sequence, with MTF2 Conferring the Strongest Affinity. (A, C, E) EMSAs examining binding between PRC2-core, PRC2–EPOP, or PRC2–MTF2 and free DNA with two Widom 601 positioning sequences separated by a 40 bp linker. Linker identities: random sequence [R; (A)], GA tandem repeats (C), or GCN tandem repeats (E). Reactions were incubated for 30 min and resolved on agarose gels; DNA was visualized by SYBR Gold staining. (B, D, F) Quantitative binding analysis for R-(B), GA-(D), and GCN-(F) linker DNA, measured from the other half of each reaction in A, C, and E, respectively, resolved on acrylamide gels for increased sensitivity. The corresponding acrylamide gels are shown in Fig. S3A–C. Data were fit with a sigmoidal binding function to calculate dissociation constants (Kd). *n* = 3 independent experiments.

**Figure S3.**
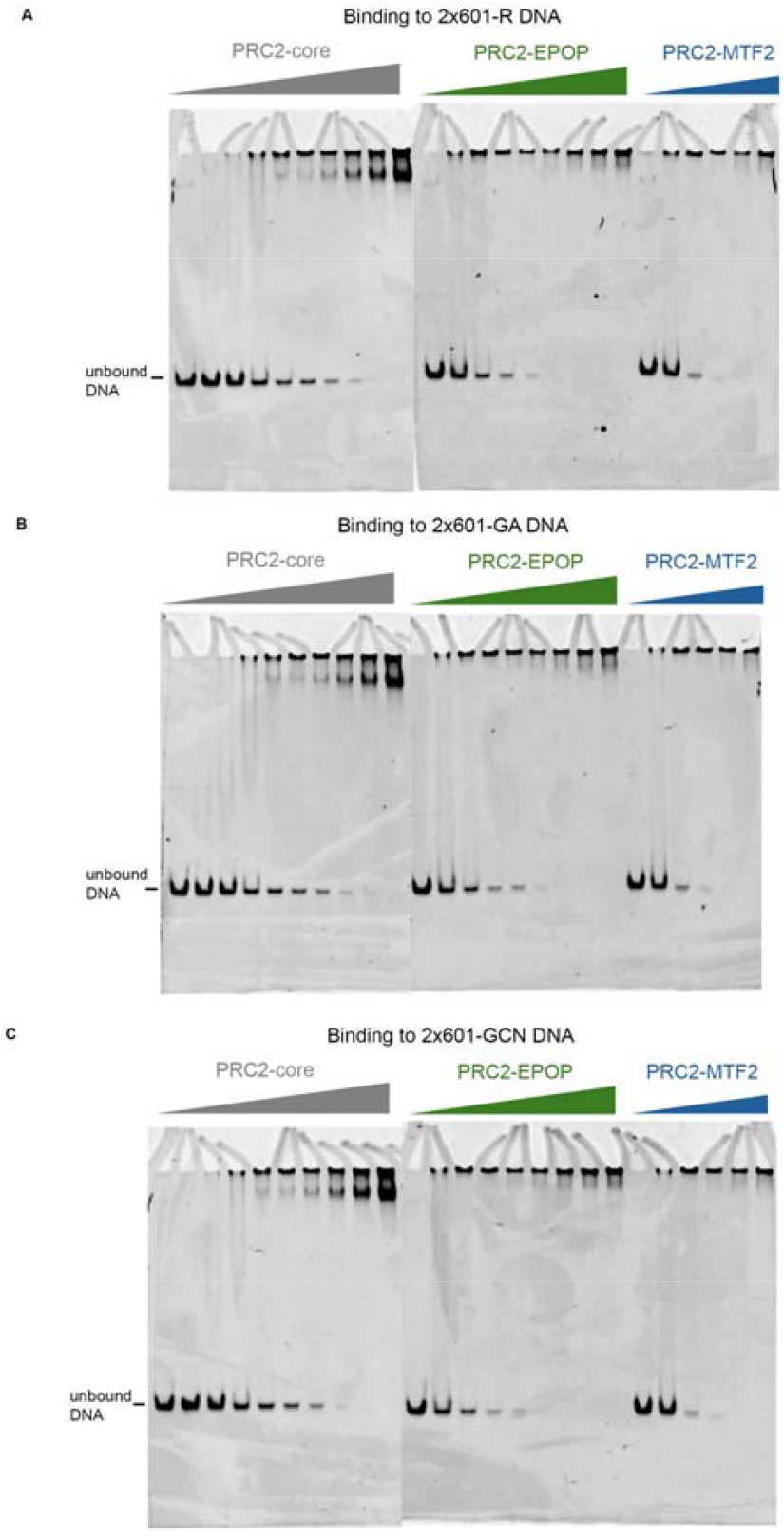
EMSAs resolved on acrylamide gels for PRC2–core, PRC2–EPOP, and PRC2–MTF2 binding to DNA with defined linker sequences, corresponding to Fig. S2. (A–C) EMSAs of PRC2–core, PRC2–EPOP, and PRC2–MTF2 incubated with free DNA containing two Widom 601 positioning sequences separated by a 40 bp linker comprising either a random sequence [R; (A)], GA tandem repeats (B), or GCN tandem repeats (C). For each condition, half of the reactions shown in Fig. S2A, C, and E were resolved on agarose gels (Fig. S2), while the other half were resolved in parallel on acrylamide gels to provide increased detection sensitivity. DNA was visualized by SYBR Gold staining. These acrylamide gels were used for quantitative analysis and curve fitting shown in Fig. S2B, D, and F.

## Notes

### Competing Interest Statement

The authors have declared no competing interest.

